# In silico Design of novel Multi-epitope recombinant Vaccine based on Coronavirus surface glycoprotein

**DOI:** 10.1101/2020.03.10.985499

**Authors:** Mandana Behbahani

## Abstract

It is of special significance to find a safe and effective vaccine against coronavirus disease 2019 (COVID-19) that can induce T cell and B cell -mediated immune responses. There is currently no vaccine to prevent COVID-19. In this project, a novel multi-epitope vaccine for COVID-19 virus based on surface glycoprotein was designed through application of bioinformatics methods. At the first, seventeen potent linear B-cell and T-cell binding epitopes from surface glycoprotein were predicted in silico, then the epitopes were joined together via different linkers. The ability of the selected epitopes to induce interferon-gamma was evaluate using IFNepitope web server. One final vaccine was constructed which composed of 398 amino acids and attached to 50S ribosomal protein L7/L12 as adjuvant. Physicochemical properties, as well as antigenicity in the proposed vaccines, were checked for defining the vaccine stability and its ability to induce cell-mediated immune responses. Three-dimensional structure of the mentioned vaccine was subjected to the molecular docking studies with MHC-I and MHC-II molecules. The results proposed that the multi-epitope vaccine with 50S ribosomal protein L7/L12 was a stable construct with high aliphatic content and high antigenicity.

## 1. Introduction

In early 2020, COVID-19 began generating headlines all over the world because of the extraordinary speed of its transmission. So, there are rising concerns about community infections. Vaccination is one of the most effective tools to prevent infectious diseases [1][2]. As far as our knowledge concerns, there is no report about developing COVID-19 multi epitope vaccine. Therefore, we became eager to design potent multiepitope vaccines from antigenic sites of coronavirus surface glycoprotein. The multiepitope vaccines have advantageous over conventional vaccines with regards to safety profile and high immunogenicity [3]. Multiepitope vaccines have the potential to induce responses restricted by a wide variety of HLA molecules and generate a balanced CD4+ and CD8+ cellular immune response. Another molecule that contribute to innate immunity contains TLR3 that activates antiviral mechanism during infection. Recently, in silico design of epitope-based vaccines has been done for vaccine developing against many infectious diseases. Some bioinformatics tools could facilitate the development of multi epitope-based vaccines. The computational tools can optimize the extensive immunological data such as antigen presentation and processing to achieve specific interpretations [4]. In recent decades, several vaccines were established based on in silico methods that include efficient vaccines against Toxoplasma gondii [5], Brucella abortus [6], Escherichia coli [7], Vibrio cholera [8], Human immunodeficiency virus-1 [9], Hepatitis C virus [10] and many others. In several experimental studies, the efficacy of computationally designed vaccines has been recently approved for use in defined human vaccines [11][13]. In this study, in silico analysis were performed to determine exclusive B cell and T-cell epitopes from coronavirus surface glycoprotein that are antigenically most significant for coronavirus. In our research, some unique exclusive B cell and T-cell epitopes from coronavirus surface glycoprotein were selected based on their antigenicity, stability and length. The selected epitopes were merged into each other using suitable linkers for organization of final vaccine construct. Consequently, the stability and efficacy of the vaccines were predicted by a set of bioinformatics methods.

## 2. Material and methods

### 2.1 Data collection

At the first step of our study, the reference amino acid sequences of coronavirus surface glycoprotein (YP-001856243.1), five HLA-1 (NP_001229971.1, NP_001229687.1, NP_002118.1, NP_061823.2, NP_005507.3) and six HLA-2 protein (NP_001229454.1, NP_006111.2, NP_001230891.1, NP_002110.1, NP_061984.2, NP_001020330.1) were retrieved from NCBI (https://www.ncbi.nlm.nih.gov). SWISS-MODEL Server (https://swissmodel.expasy.org/) was utilized for modelling of 3-D structures of HLA class I and HLA class II, But for TLR-3 the data in PDB bank was used and optimized by chimera 1.12 [14.]

### 2.2 Multiple sequence alignment and antigen selection

To determine exclusive conserved sequence of the coronavirus surface glycoprotein, NCBI BLAST was performed (https://blast.ncbi.nlm.nih.gov/Blast.cgi). Also, for defining the conserved region (s) in the protein sequences, multiple sequence alignment was done by Multalin server (https://www.multalin.toulouse.inra.fr/multalin). Additionally, the antigenicity of the coronavirus surface glycoprotein was evaluated using VaxiJen 2.0 server (http://www.ddg-pharmfac.net/vaxijen/VaxiJen/VaxiJen.html[15]. Finally, the most particular conserved sequence and antigenic peptide were selected for further analysis.

### 2.3 B-cell epitope prediction and selection

Linear B-cell epitopes in the vaccine model were predicted using ElliPro (http://crdd.osdd.net/raghava/bcepred) [16] and IEDB analysis Resource (http://tools.iedb.org/population) [17].

### 2.4 T-cell epitope prediction and selection

MHC-I restricted epitopes were predicted through ProPred-1 server (http://tools.immuneepitope.org/analyze/html/mhc_binding.html) [18]. The server uses special patterns for HLA-A*03:01 allele. Similarly, MHC-II restricted epitopes were predicted using ProPred server (http://tools.immuneepitope.org/mhcii). The server uses special patterns for DRB1*01:07 allele. Finally, the conserved sequence and antigenic peptide were selected for further analysis.

### 2.5 Construction of final vaccine

Seventeen suitable common B-cell and T-cell epitopes (9-16 amino acids) from coronavirus surface glycoprotein were selected and organized in the final vaccine construct. Then, these epitopes were merged together with AAY, KK linkers and considered as a multi-epitope vaccine. One adjuvant “50S ribosomal protein L7/L12 (MSDINKLAETLVNLKIVEVNDLAKILKEKYGLDPSANLAIPSLPKAEILDKSKEKTSFDLILKGAGSAKLTVVKRIKDLIGLGLKESKDLVDNVPKHLKKGLSKEEAESLKKQLEEVGAEVELK)

with 124 amino acids were incorporated with **EAAAK** linker at N-terminal portion of the constructs. The sequence of the designed vaccine structures with their adjuvant are depicted in Table 3. The final vaccines (IV1) stretch was found to be 398 amino acids.

### 2.6 Physicochemical properties analysis

In this research, five characteristics (molecular weight, theoretical pI, extinction coefficient, aliphatic index and grand average of hydropathicity) of the constructed vaccine was evaluated using ProtParam server (http://web.expasy.org/protparam).

### 2.7 Secondary structure analysis

The frequency of the secondary structure of the constructed vaccines (alpha helix, extended strand and random coil) were computed using GOR IV web server (http://npsa-pbil.ibcp.fr/cgi-bin/npsa_automat.pl?page=npsa_gor4.html).

### 2.8 Molecular docking study

To confirm the binding affinity of the best vaccine to MHC-I and MHC-II molecules, molecular docking was done between the selected vaccines and five HLA-1 structures (NP_001229971.1, NP_001229687.1, NP_002118.1, NP_061823.2, NP_005507.3), also six HLA-2 proteins (accession numbers: NP_001229454.1, NP_006111.2, NP_001230891.1, NP_002110.1, NP_061984.2, NP_001020330.1) and TLR-3 (PDB ID: 2A0Z) separately. Molecular docking studies were done using H-dock server (https://bioinfo3d.cs.tau.ac.il/PatchDock/) with default complex type and clustering RMSD of 4Å. The binding sites of final construct and TLR3 were studied using fully automated protein-ligand interaction profiler server (PLIP; https://projects.biotec.tu-dresden.de/plip-web/plip/index). The outputs of PLIP were in XML format, flat text, and visualization files [19]. The visualization files were visualized using PyMOL software (windows version 2.0.7). Then, seventeen epitopes were analysed by IFNepitope server.

### 2.10 IFNγ analysis

IFNepitope web server is established for users working in the field of vaccine design. This server allows users to predict and design IFN-gamma inducing peptides. The ability of the selected epitopes to induce interferon-gamma was evaluate using IFN epitope server.

## 3. Results

### 3.1 Multiple sequence alignment and antigen selection

Initially, coronavirus surface glycoprotein was studied for determining specific conserved part of protein between the virus serotypes. Results of protein BLAST are shown in Table 1. The results demonstrated that coronavirus surface glycoprotein had the most conservancy levels between 97. 8 and 100. Also, VaxiJen score of the protein showed high antigenicity. Due to having good antigenicity, high exposure probability to the immune system and high conservancy, this protein was selected for vaccine design.

**Table 1:**
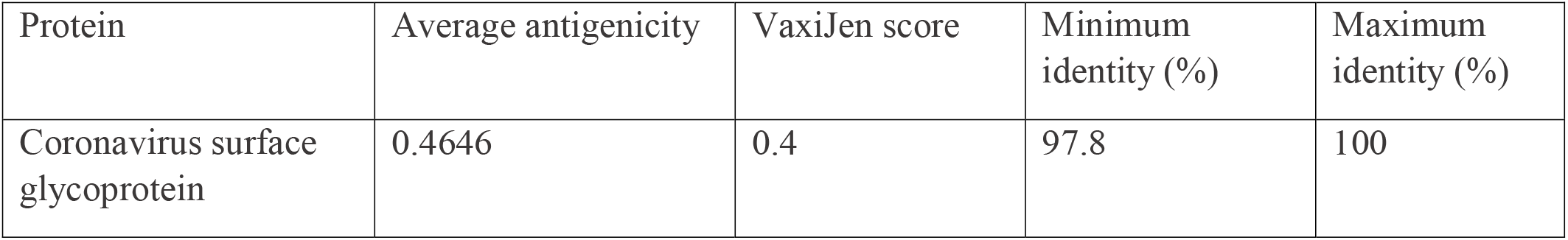
Results of the antigenicity and BLAST of coronavirus surface glycoprotein and antigenicity prediction

### 3.2 T-cell and B-cell epitope prediction

In the current study, some appropriate common B-cell and T-cell epitopes were designed. The predicted MHC-I and MHC-II restricted epitopes were compared to B-cell epitopes to determine shared epitopes (Table 2). Finally, 17 epitopes with 9-16 amino acids were selected. These epitopes are located between residues 14-642. These epitopes exhibited a relatively high aliphatic index (>60), high antigenicity (>1.6) and low instability index (less than 20). The most potent epitope was repeated three times and merged into the other epitopes using suitable linkers (KK, AYY) for organization of final vaccine construct.

**Table 2:**
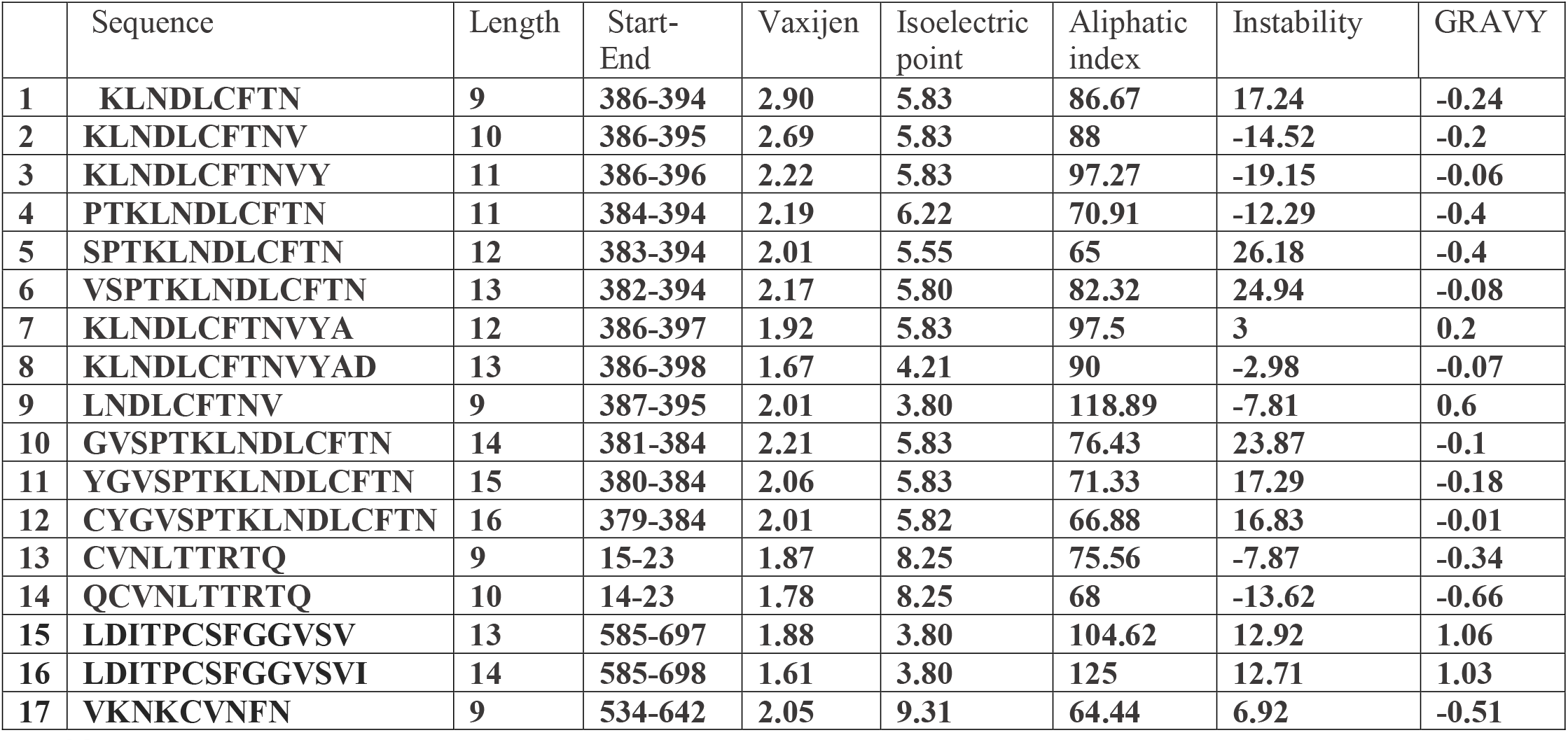
Result of final T-cell and B-cell epitope prediction screening from coronavirus surface glycoprotein

### 3.3 Antigen selectivity of constructed vaccines

Final construct vaccine was composed of 398 amino acids which were respectively attached to 50S ribosomal protein L7/L12 as adjuvant. The antigenicity score of constructed vaccine are shown in Table 3. The result demonstrated that IV1 has 1.2110 antigenicity (Table 3).

**Table 3:**
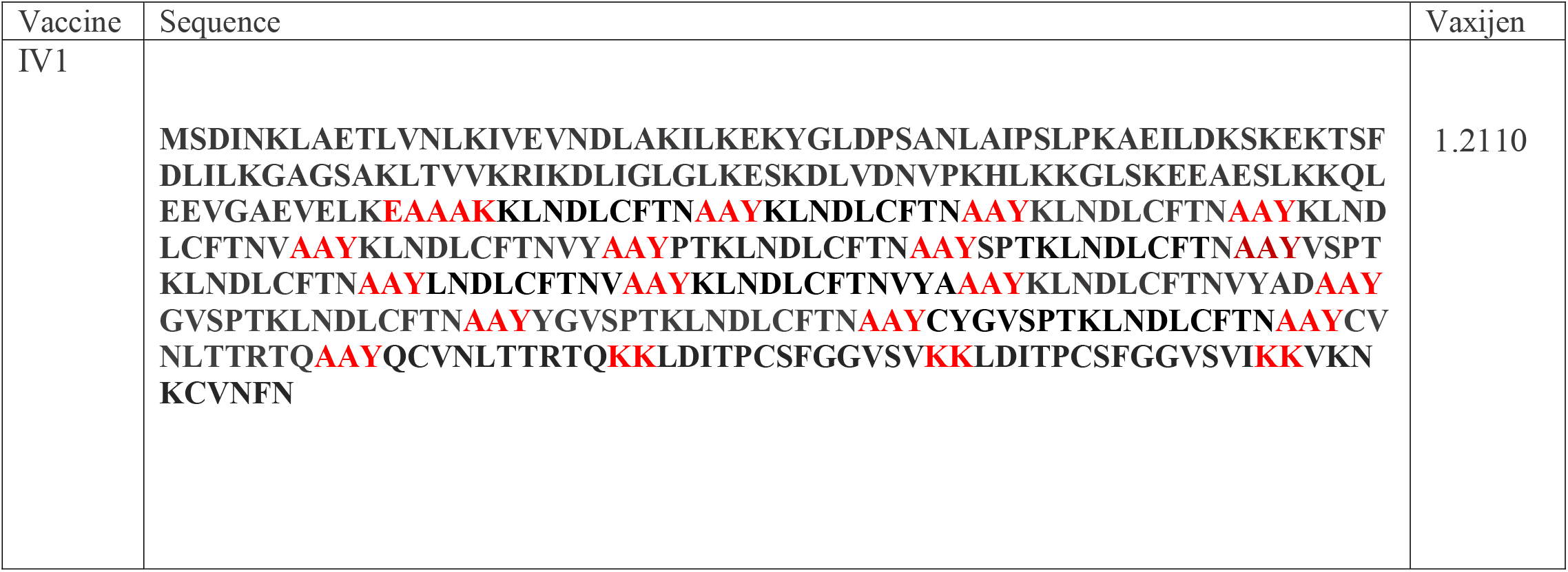
Average antigenicity of constructed vaccine using Vaxijen

### 3.4 Physicochemical properties

Physicochemical properties of the constructed vaccine were predicted using Protparam server. The results revealed that this multi-epitope vaccine have low instability “as a value below 40” predicts that the protein is stable. The IV1 construct showed the highest Isoelectric point with 8.52 value. From the aliphatic Index of view, this construct showed aliphatic index more than 90 % (Table 4). The results of gor4 demonstrated that the random coil values of the IV1 were high compared to Alpha helix and extended structure.

**Table 4:** Physico-properties of constructed vaccines

| Protein | Molecular weight | Isoelectric point | Aliphatic index | GRAVY | Instability | Alpha | Extended | Random coil |
| --- | --- | --- | --- | --- | --- | --- | --- | --- |
| IV1 | 43679.43 | 8.52 | 92.71 | -0.046 | 11.85 | 23.37% | 8.54% | 68.09 |

### 3.5 Analysis of docking results

The results of docking the constructions with HLA-1 and HLA-2 confirmed the high values with IV1 construct. Also, the results of TLR-3 analysis showed values of 1680.569 for IV1 which demonstrate the high efficacy of vaccine construct (Tables 5). Figure 2 indicate docking results of the Vac1 construct with the TLR-3 as the examples of vaccines potential for interaction. The results showed that TLR-3 have interaction to Glu115, Gly 118 residues of AV1.

**Fig 1:**
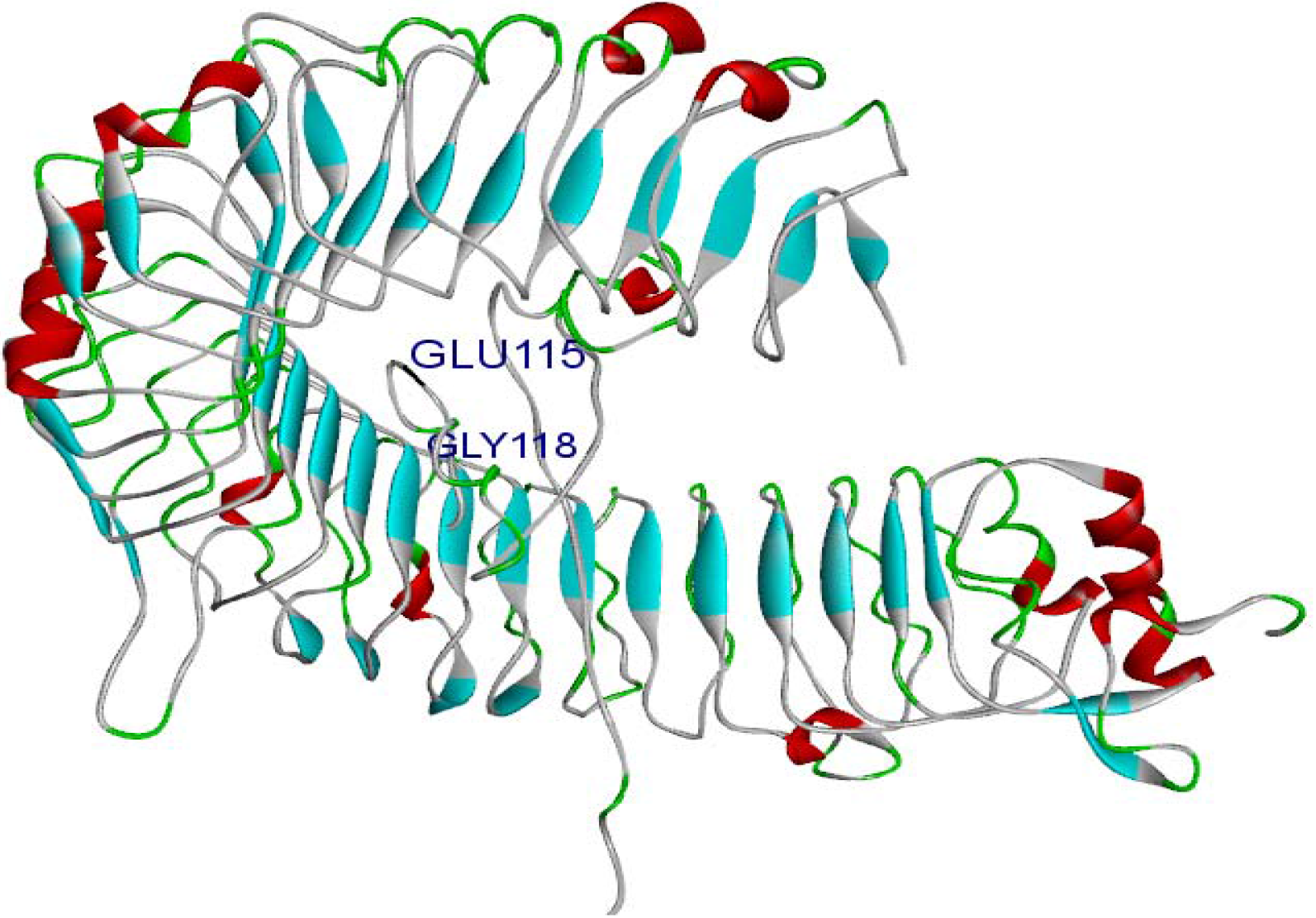
Molecular docking analysis of IV1construct with TLR-3. The results showed that TLR-3 have interaction to Glu 115 and Gly 118 residues of AV1

**Table 5:** HDOCK scores of HLA1, HLA2 and interaction residues of selected sequences

| Protein | Hla1-A1 | Hla1-Chain G | Hla1-Chain E | Hla1-Chain F | Hla1-Chain CW-1 | TLR3 | Hla2 - antigen gama chain | Hla2-DM | Hla2-DO | Hla2-DP | Hla2-DQ | Hla2- DR |
| --- | --- | --- | --- | --- | --- | --- | --- | --- | --- | --- | --- | --- |
| IV1 | 951.2 | 957.2 | 951.2 | 1215.7 | 1220.8 | 1680.5 | 1215 | 1214 | 1215.7 | 1215.7 | 1215.7 | 1215.7 |

### 2.10 IFNγ analysis

The result of IFNγ analysis was described in Table 6. The result demonstrates that among 17 epitopes, 16 epitopes have potential to produce IFN-γ. Epitope 6 showed the highest score with value of 2.

**Table 6:**
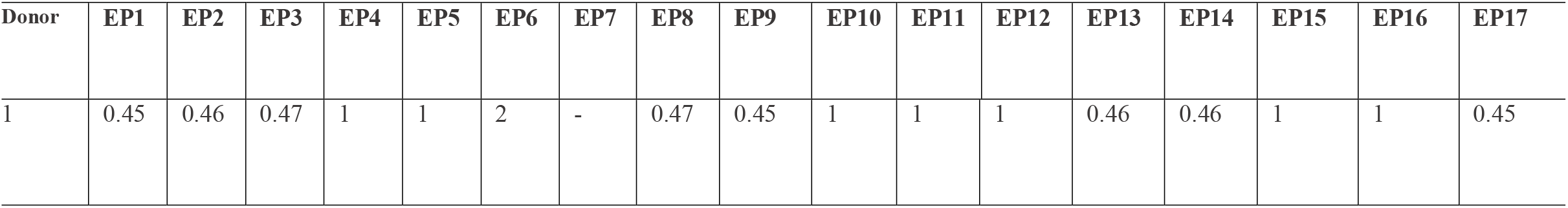
IFNγ analysis score of 17 selected epitopes using IFNepitope web server

## 4. Discussion

Due to the nature of coronavirus and high infectious rate, the progress of a vaccine against coronavirus is very challenging. Though, with the development of computational methods, these limitations are reduced. By the way, using computational methods, the design of recombinant vaccines and the estimation of physicochemical properties as well as vaccines efficacy could be available [20] [21]. Then, this research was intended to design an effective multi-epitope recombinant vaccine against coronavirus using a unique multi-step bioinformatics approach. Our potent multi-epitope vaccine is contained seventeen epitopes in surface glycoprotein of coronavirus along with AAY and KK linkers and 50S ribosomal protein L7/L12 as adjuvant. Based on our knowledge, there is no report about computational design of epitope-based vaccine for coronavirus. Recently, in silico, design of epitope-based vaccine was used for vaccine development against several infectious diseases. Several bioinformatics tools have been established that accelerate the growth of multi epitope-based vaccines. In recent decade, several multiepitope vaccines for pathogenic viruses have been reported. Multi epitope vaccines could provide an effective immunization against different serotypes of a pathogen. Despite mentioned advantages of Multi epitope vaccines, poor immunogenicity is considered as a major drawback to growth of these vaccines [22], [23]. The in-silico results proposed that our multi-epitope vaccine was very stable with high aliphatic index and it was potentially antigenic. As reported earlier, high aliphatic index shows the higher thermos-stability of the constructed vaccine. At the present research, the aliphatic index was high, and instability was low and Gravy indices were negative. Also, the higher PI, as the case of this study, shows the higher potential for cell wall attachment. However, the proposed vaccine has high antigenicity. This was chosen for docking studies. The results demonstrated that our mentioned vaccine could be a right candidate for experimental research [24].

## Conclusion

This study introduced designing novel multi epitope vaccines against Coronavirus which could cover conserve sequence of the virus. The multi-epitope vaccine presented by this study showed promising result through in silico step, which could be followed by in vitro and in vivo studies.

## Conflict of interest

Authors declare no conflict of interest.

## Acknowledgment

We wish to thank the University of Isfahan for their supports.

## Notes

#### Summary of Updates

some results in new version has been changed

## References

1. Li Q., Guan X., Wu P., Wang X., Zhou L (2020) Tong Y., Feng Z. Early transmission dynamics in Wuhan, China, of novel coronavirus-infected pneumonia. New England Journal of Medicine.

2. World Health Organization Novel coronavirus (2019-nCoV) situation reports. 2020. https://www.who.int/emergencies/diseases/novel-coronavirus-2019/situation-reports

3. Cicala C, Nawaz F, Jelicic K, et al. (2016) HIV-1 gp120: A Target for Therapeutics and Vaccine Design. Current Drug Targets. 17(1):122–35.

4. Kumar M, Thakur V, Raghava GP (2008) “COPid: composition based protein identification. In silico b10.Soria-Guerra RE, Nieto-Gomez R, Govea-Alonso DO, et al. An overview of bioinformatics tools for epitope prediction: implications on vaccine Development. J Biomed Inform. 2015;53, 405–414. doi: 10.1016/j.jbi.2014.11.003.iology. 8:121–128.

5. Soria-Guerra RE, Nieto-Gomez R, Govea-Alonso DO (2015) t al. An overview of bioinformatics tools for epitope prediction: implications on vaccine Development. J Biomed Inform. 2015;53, 405–414. doi: 10.1016/j.jbi.2014.11.003.

6. Bissati KE, Chentoufi AA, Krishack PA (2016) Adjuvanted multi-epitope vaccines protect HLA-A* 11: 01 transgenic mice against Toxoplasma gondii. JCI Insight. 1(15): doi: 10.1172/jci.insight.85955.

7. Escalona E, Saez D, Onate A (2017) Immunogenicity of a multi-epitope dna vaccine encoding epitopes from Cu–Zn superoxide dismutase and open reading Frames of Brucella abortus in mice. Frontiers Immunology. 8, 125

8. Rodrigues-da-Silva RN, Martins da Silva JH, Singh B (2016) In silico identification and validation of a linear and naturally immunogenic B-cell epitope of the Plasmodium vivax malaria vaccine candidate merozoite surface protein-9. Plos One;11(1): e0146951. doi: 10.1371

9. Nezafat N, Karimi Z, Eslami M (2016) Designing an efficient multi-epitope peptide vaccine against Vibrio cholerae via combined immunoinformatics and protein interaction based approaches. Computational Biology and Chemistry. 62: 82–95. doi: 10.1016/j.compbiolchem.2016.04.006

10. Yang Y, Sun W, Guo J, et al. In silico design of a DNA based HIV-1 multi-epitope vaccine for Chinese populations. Human Vaccines Immunother. 2015;11(3): 795–805. doi: 10.1080/21645515.2015.1012017

11. Nosrati M, Mohabatkar H, Behbahani M (2017) A novel multi-epitope vaccine for cross protection against Hepatitis C Virus (HCV): an immunoinformatics. Approach Research in Molecular Medicine 5(1): 17–26.

12. Rahjerdi AK, Amani J, Rad I (2016) Designing and structure evaluation of multi-epitope vaccine against ETEC and EHEC, an in silico approach. Protein Peptide Letters. 23(1): 33–42.

13. Oscherwitz J (2016) The promise and challenge of epitope-focused vaccines. Human Vaccines Immunother.; 12(8): 2113—2116.

14. Conrad C. Huang, Elaine C. Meng, John H. Morris (2014) Enhancing UCSF Chimera through web services. Nucleic Acids Research. 42(1):478–484. https://doi.org/10.1093/nar/gku377

15. Doytchinova IA, Flower DR (2007) VaxiJen: a server for prediction of protective antigens, tumour antigens and subunit vaccines. BMC Bioinformatics.

16. Duquesnoy R, Marrari M (2017) Usefulness of the ElliPro epitope predictor program in defining the repertoire of HLA-ABC eplets. Hum. 78(7-8): 481–488. doi:10.1016/j.humimm.03.005

17. Beaver JE, Bourne PE, Ponomarenko JV (2007) Epitope Viewer: a Java application for the visualization and analysis of immune epitopes in the Immune Epitope Database and Analysis Resource (IEDB). Immunome Res DOI:10.1186/1745-7580-3-3.

18. Patronov AI. Doytchinov I (2013) T-cell epitope vaccine design by immunoinformatics. Open Biol. 2013; 3(1):120139. doi: 10.1098/rsob.120139.

19. Salentin S, Schreiber S, Haupt VJ (2015) PLIP: fullyautomated protein–ligand interaction profiler. Nucleic Acids Research; 43: W443–W447. https://doi.org/10.1093/nar/gkv315

20. Roosa K., Lee Y., Luo R., Kirpich A., Rothenberg R., et al (2020) Hyman J.M., Yan P., and G. Chowell, Real-time forecasts of the COVID-19 epidemic in China from February 5th to February 24th, 2020, Infect Dis Model. 2020; 5: 256–263.

21. Ai S., Zhu G., Tian F., Li H., Gao Y., Wu Y., Lin H (2020) Population movement, city closure and spatial transmission of the 2019-nCoV infection in China.

22. Paul S, Piontkivska H (2010) Frequent associations between CTL and T-Helper epitopes in HIV-1 genomes and implications for multi-epitope vaccine designs. BMC Microbiology. 10:212. doi: 10.1186/1471-2180-10-212

23. Hajighahramani N, Nezafat N, Eslami M, et al. (2017) Immunoinformatics analysis and in silico designing of a novel multiepitope peptide vaccine against Staphylococcus aureus. Infection Genetics and Evolution. 48: 83–94. doi: 10.1016/j.meegid

24. Solanki V, Tiwari M, Tiwari V. (2019) Prioritization of potential vaccine targets using comparative proteomics and designing of the chimeric multi-epitope vaccine against Pseudomonas aeruginosa. Scientific Reports. 9.

